# Computational evidence of a compound with nicotinic α4β2-Ach receptor partial agonist properties as possible coadjuvant for the treatment of obesity

**DOI:** 10.1101/088138

**Authors:** Helena den-Haan, Juan José Hernández Morante, Horacio Pérez-Sánchez

**Author notes:** Corresponding authors: Horacio Pérez-Sánchez, And Juan José Hernández Morante.

## Abstract

**Background:** Nowadays, the search for new anti-obesity drugs is oriented to the use of anti-addiction medications like bupropion and naltrexone. Other compounds like varenicline may be also useful to treat obesity. However, the low effectiveness of the former or the high number of adverse effects of the latter makes it necessary to search for new therapeutic agents.

**Methods:** Screening database selected for the computational experiments was DrugBank. 3D global shape comparison with varenicline was performed by means of the Ligand Based Virtual Screening tool WEGA v2015. A pharmacophore model based in the structure of varenicline was created by means of LigandScout v4.08. The in-silico screening was performed using Relative Pharmacophore Fit (RPF) scoring function implemented in LigandScout. Up to 3 mismatches with varenicline pharmacophore model were allowed for hits retrieving.

**Results:** Drugbank database was screened in silico to find alternative molecules to varenicline, and the compound cevimeline was found to have strong similarity to varenicline in terms of 3D shape and pharmacophoric features. Thus, we propose this hit may interact with nicotinic α4β2-Ach receptor in the same mode as varenicline does.

**Discussion:** The functional activities of this compound and its validity as a drug therapy for obesity treatment must be confirmed in further in vitro, in vivo and preclinical studies; however, attending to our screening procedure, this compound should be a promising therapy for such a complex disorder such as obesity.

## 1. INTRODUCTION

In most countries, overweight and obesity are still the most prevalent health problems, affecting more than 50% percent of the population (more than 1 in 2 subjects) (WHO). As overweight and obesity progress, the presence of certain related-diseases such as diabetes, cardiovascular disease and cancer also increases (Bosello, Donataccio & Cuzzolaro, 2016).

This situation highlights the need to address overweight and obesity through effective therapeutic strategies in order to achieve significant weight loss in the short term, which should be maintained over time. This is a key issue, because as it has been verified in a recent systematic review, the effectiveness of low-calorie diet interventions is modest, especially in the long term (Langeveld & DeVries, 2015). In addition, according to that study, the variability among subjects regarding the effectiveness of treatment was very broad, so in general, these therapies cannot achieve the therapeutic objectives in overweight patients (Langeveld & DeVries, 2015).

Therefore, the interests of researches are directed towards the development of new therapeutic interventions for the treatment of overweight. Specifically, drug therapies have been developed mainly focused in the inhibition of fat absorption, decreasing the total amount of calorie intake; however, this blockade of fat absorption produce a huge number of adverse gastrointestinal effects that limit the effectiveness of these intervention (Douglas et al., 2015).

Consequently, a great advance into the molecular basis of appetite regulation has been performed to identify new therapeutic targets. Hence, a key advance came from the demonstration of the importance of the “reward center”, a brain region located in the nucleus accumbens (NAc) associated with certain addictions such as tobacco and opioids, in appetite regulation (Kenny, 2011).

In this regards, several studies have observed that sucrose is able to active dopamine release at the “reward centre” through the activation of nicotinic α4β2-Ach receptor, in the same way that certain substances like tobacco (Anderberg et al., 2014).

A previous study in rats showed that intense sweetness commonly seen in sugar-rich diets tends to override self-control mechanisms in the body by modifying the “reward centre” in the brain, hence increasing susceptibility to highly palatable foods (Lenoir et al., 2007). Although unproven in humans, this evidence could serve as a starting point for an early reassessment of initiatives designed to control the excessive consumption of sugar at the present time.

For these reason, the new strategies for obesity treatment are focused on the use of new drugs initially developed to the treatment of drug addictions, like bupropion, a dopamine reuptake inhibitor (Yanovski & Yanovski, 2015). Nevertheless, this treatment does not appear to be effective in the long term either. Moreover, other smoking cessation drugs like varenicline, a partial agonist of nicotinic α4β2-Ach receptors, seem to be more effective that bupropion, but with more adverse effects (Cinciripini et al., 2013). Therefore, although the use of drugs for addiction seems to be a good strategy for obesity treatment, it is still necessary to develop new effective and with fewer side effects drugs for patients with overweight.

At this moment, the development of new drugs has also progressed enormously, mainly with the incorporation of structural bioinformatics, computational biochemistry and super-computational techniques, which as a whole, through a virtual molecular modeling allow us to identify “target” molecules with the desired therapeutic activity. Other approach to drug discovery may be through the use of known molecules, which will allow discovering other molecules that will share similar pharmacological properties, especially regarding molecule-receptor interactions (Merelli et al., 2014).

For all the commented, the present work was conducted to discover or identify molecules that interact with nicotinic α4β2-Ach receptors in a similar way that it does varenicline, sharing a similar three-dimensional shape and pharmacophoric properties with varenicline.

Based on these computational predictions, it is hypothesized that the identified molecule should be effective for the obesity treatment through the inhibition of the hedonic response of palatable foods, which may help patients to control food intake and especially to avoid relapse, a fact that may increase the effectiveness of obesity treatment but especially at long-term, precisely the weakness of current treatments.

## 2. MATERIAL & METHODS

Screening database selected for the experiments was DrugBank (Wishart et al., 2006), approved drugs subset, retrieved in November 2015. Up to 200 conformations were generated for every molecule with omega (Hawkins et al., 2010). The other parameters were set to default. 3D global shape comparison with varenicline was performed by means of WEGA v2015 using pure shape scoring function (Yan et al., 2013). To avoid excluding possible hits a threshold score of 0.5 was set. Thus, all molecules in Drugbank approved database subset scoring more than 0.5 were kept. A pharmacophore model based in the structure of varenicline was created by means of LigandScout v4.08 (Wolber & Langer, 2005) (Fig 1). Screening database was prepared in LigandScout by means of idbgen omega-best parameters. The screening was performed using Relative Pharmacophore Fit (RPF) scoring function implemented in LigandScout. Up to 3 mismatches with varenicline pharmacophore model were allowed for hits retrieving. Scoring functions from WEGA and RPF score the amount of similarity and feature matching respectively and normalize that value to a 0-1 scale.

**Figure 1.**
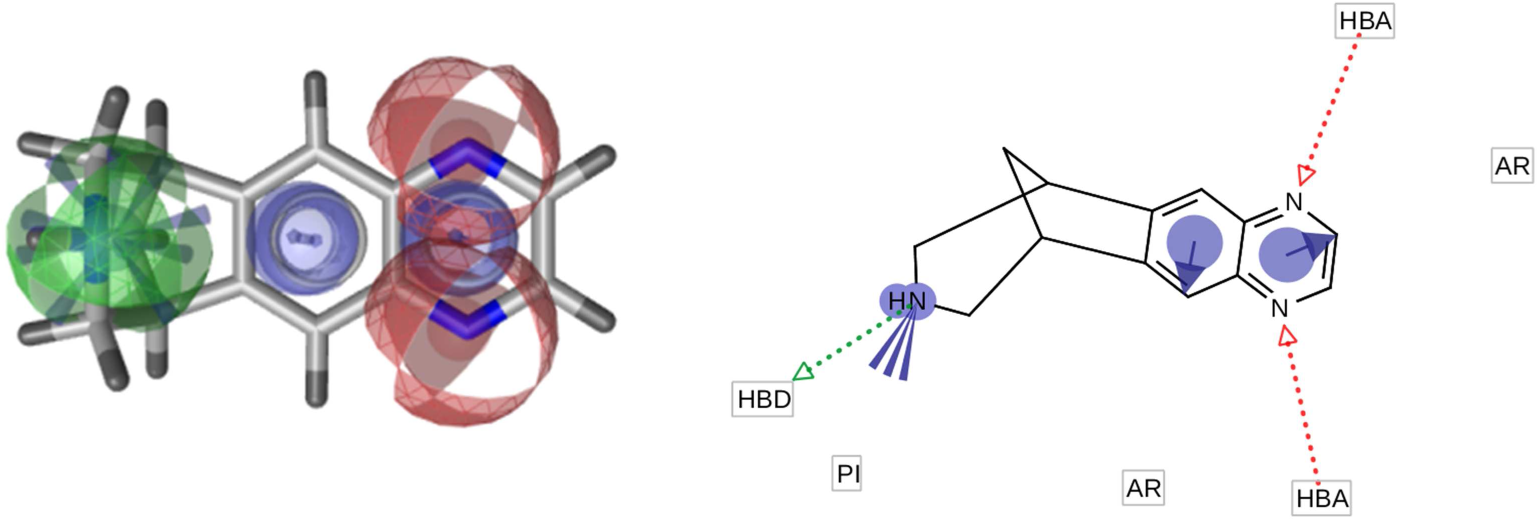
Pharamacophore model of varenicline. Features in the model are: 1 Positive Ionizable group (blue star), 1 Hydrogen Bond Donor (green sphere), 2 Aromatic Rings (blue rings), 2 Hydrogen Bond Acceptors (red spheres). From left to right 3D and 2D figures are presented from the pharmacophore model overlapped on Varenicline.

## 3. RESULTS

After having screened all approved drugs in DrugBank by means of WEGA and LigandScout a total of 288 compounds were found to have a 3D shape similarity score of 0.5 or higher and fit the varenicline pharmacophore model with a Relative Pharmacophore Fit score ranging from 0.64 to 0.94. Seven of these predicted compounds were of special interest since their scores ranged from ~0.80-0.86 in shape similarity score and at the same time they showed a predicted pharmacophore fitting of ~0.9-0.94 (Table 1) as considered a more consistent prediction. Known protein targets for every hit are enumerated, along to a 2D diagram of the retrieved drug showing the pharmacophore features common to varenicline.

**Table 1.**
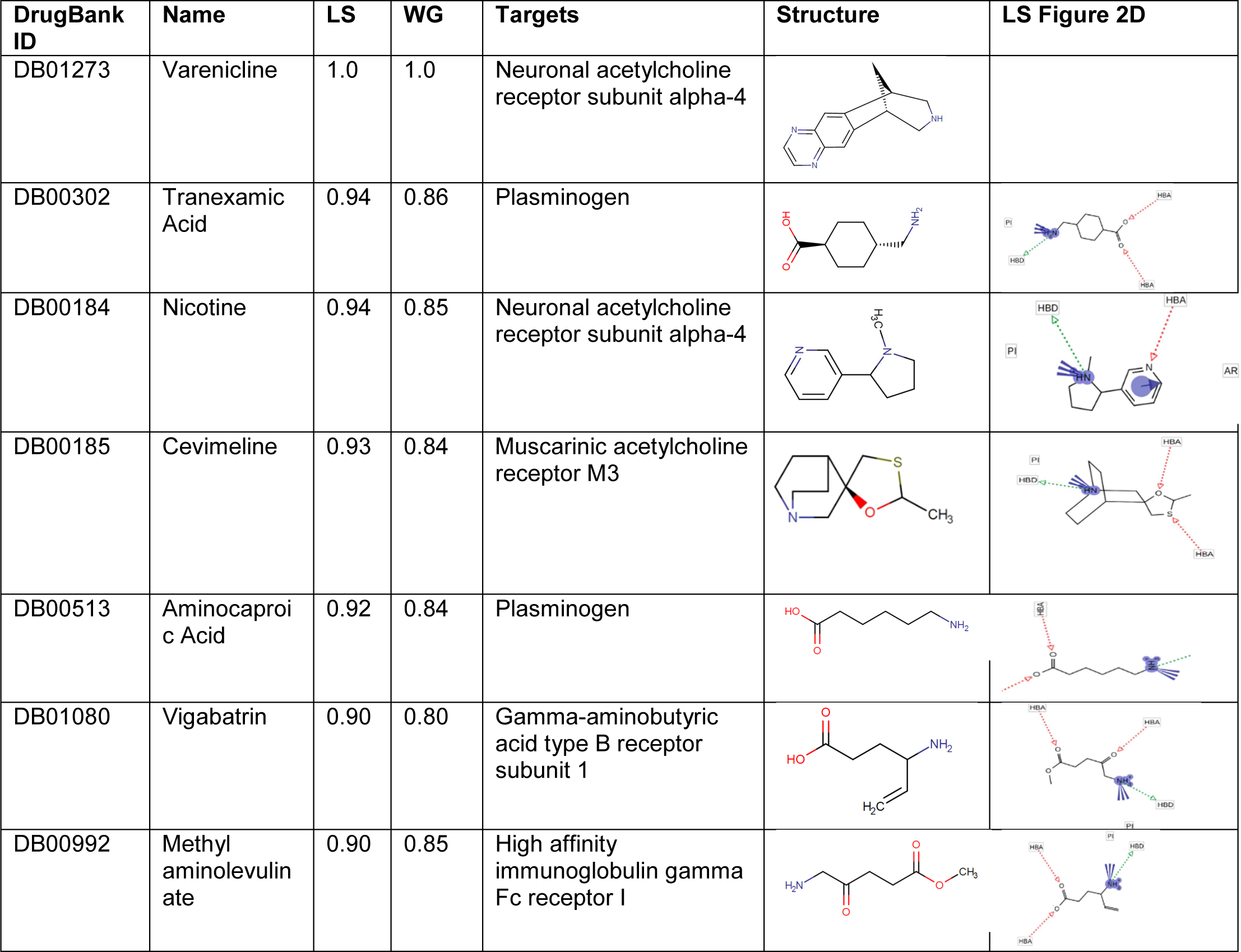
List of compounds included in Drugbank database for which high 3D shape similarity (WG) and alignment score of their respective pharmacophoric maps (LS) to varenicline where calculated.

| DrugBank ID | Name | LS | WG | Targets | Structure | LS Figure 2D |
| --- | --- | --- | --- | --- | --- | --- |
| DB01273 | Varenicline | 1.0 | 1.0 | Neuronal acetylcholine receptor subunit alpha-4 |  |  |
| DB00302 | Tranexamic Acid | 0.94 | 0.86 | Plasminogen |  |  |
| DB00184 | Nicotine | 0.94 | 0.85 | Neuronal acetylcholine receptor subunit alpha-4 |  |  |
| DB00185 | Cevimeline | 0.93 | 0.84 | Muscarinic acetylcholine receptor M3 |  |  |
| DB00513 | Aminocaproic Acid | 0.92 | 0.84 | Plasminogen |  |  |
| DB01080 | Vigabatrin | 0.90 | 0.80 | Gamma-aminobutyric acid type B receptor subunit 1 |  |  |
| DB00992 | Methyl aminolevulinate | 0.90 | 0.85 | High affinity immunoglobulin gamma Fc receptor I |  |  |

## 4. DISCUSSION

The present work was developed in order to find molecules analog to varenicline, with a similar partial agonist activity on nicotinic α4β2-Ach receptors. After screening the approve drugs subset of the Drugbank database, our data retrieve cevimeline, a murscarinic receptor agonist, like the most suitable molecule that should be able to interact with this receptor in a similar way than varenicline does.

Cevimeline is indicated for the treatment of symptoms of dry mouth in patients with Sjögren’s Syndrome. Over the central nervous system, cevimeline act as a M1 agonist, but its affinity is greater on M3 receptors from epithelial salivary and lacrimal glands. As a stimulant of muscarinic receptors in general, cevimeline stimulates the secretion of the exocrine glands (Ono et al., 2004).

This affinity for several different acetylcholine receptors has been observed for several molecules. For instance, nicotin is able to interact with diverse conformations of acetylcholine receptor populations, both nicotinic and muscarinic. Pilocarpine and scopolamine also are able to interact with these receptor as selective agonist or antagonist (Nunes et al., 2013).

These data as a whole reinforce the need to repurpose the therapeutic benefits of cevimeline. In the revised bibliography, there are different examples of medications firstly developed to an end and finally used to other, like sildenafil, which was firstly develop to the treatment of erectile disfunction but it has been revealed like a promising agent against pulmonary hypertension (Vasquez et al., 2016). Therefore, in the case that our observations were confirmed, we will establish, for the first time, the novel therapeutic activity of cevimeline.

To date, the search for alpha4beta2 receptors have produced a high amount of different drugs with pleiotropic effects. For this reason, we focused on varenicline, a drug able to both partially stimulate nicotinic α4β2-Ach receptors in the absence of nicotine, while preventing nicotine from triggering a full response (Hone et al., 2016). Since its role on smoking cessation has been well established, there are very few studies focused on its role on sugar-addiction, and only a previous study indirectly evaluated its role on body weight regulation (Mangubat et al., 2012).

Furthermore, the use of drugs like bupropion or naltrexone, firstly developed to treat drug addictions, for the treatment of obesity, confirm, at least in part, the usefulness of addictive-cessation drugs. Nevertheless, the effectiveness of these drugs is limited, and the appearance of major adverse cardiovascular events in obese patients with cardiovascular risk factors has brought some controversy about the cardiovascular safety of these anti-obesity agents (Nissen et al., 2016). On the other hand, varenicline has numerous adverse effects with commonly are related with treatment discontinuation (Drovandi, Chen & Glass, 2016), and even a case of delirium have been previously reported (Virupaksha et al., 2015).

There are several reasons that could be confirming the effectiveness of our approach. On the one hand, our screening revealed nicotine, the natural ligand of nicotinic α4β2-Ach receptors. On the other hand, the drug vigabatrin, which was also suggested in our analysis, has been previously proposed as an anti-obesity agent (DeMarco et al., 2008). Interestingly, this irreversible inhibitor of GABA transaminase was also described as a potent agent against cocaine sensitization. (Gardner et al., 2002). Therefore, it is tempting to speculate that the compound selected in our in-silico screening procedure, cevimeline, could perform the same functions, at least in reference to their anti-obesity properties.

Our ligand based comparison revealed other remarkable compounds. In this regard, the pro-thrombotic drugs tranexamic acid and aminocaproic acid were found to be highly similar to varenicline. However, taking into account that one of the key features related to obesity is the elevated risk of thrombosis and cardiovascular disease, the use of these drugs could be more harmful than beneficial. Even so, our data revealed an interesting field that should be explored in further studies.

The limitations of this study should be also discussed. Evidently, the most important limitation of this study is its own design; since this is a preliminary screen of compounds with similar pharmacological properties that varenicline. Therefore, the interactions observed in silico should be validated at least in vitro. In this regard, the use of in vitro electrophyisiological studies is mandatory. This should be the first step of a long race to the final development of a possible drug for obesity treatment.

## 5. CONCLUSION

In summary, in the present work we propose that cevimeline, with a predicted partial agonism activity on nicotinic α4β2-Ach receptors, shares similar pharmacological properties with varenicline. Attending to the role of this drug, targeting nicotinic α4β2-Ach receptors with cevimeline as a coadjuvant factor would provide improve efficacy in the treatment of obesity, reducing craving and withdrawal symptoms in obese patients undergoing a balanced hypocaloric dietary treatment.

## ACKNOWLEDGEMENTS

This work was partially supported by the Fundación Séneca del Centro de Coordinación de la Investigación de la Región de Murcia under Project 18946/JLI/13. This research was partially supported by the supercomputing infrastructure of Poznan Supercomputing Center, and by the e-infrastructure program of the Research Council of Norway, and the supercomputer center of UiT - the Arctic University of Norway. This work was also funded by the PMAFI Project 06/12 of the Catholic Univesity of Murcia Research Program.

